# Identification of a second site for coat protein binding in bacteriophage P22 scaffolding protein

**DOI:** 10.1101/2022.12.13.520362

**Authors:** Corynne L. Dedeo, Richard D. Whitehead, Carolyn M. Teschke

## Abstract

Scaffolding proteins are essential for the assembly of most tailed, double-stranded DNA bacteriophages as well as herpesviruses. These proteins interact specifically with the coat proteins to efficiently assemble procapsids with the correct morphology. A helix-turn-helix (HTH) domain of bacteriophage P22 scaffolding protein is essential for coat binding, but the presence of additional coat protein binding sites has been predicted. An alanine substitution at scaffolding protein residue L245 causes a strong cold-sensitive phenotype. Both *in vivo* and *in vitro* assembly with L245A scaffolding protein yields aberrant and petite particles at non-permissive temperatures. The L245A scaffolding protein is destabilized as determined by thermal melts monitored by circular dichroism. Through crosslinking studies, residue L245 interacts with the coat protein A-domain residue D198, which has been predicted previously to contain a scaffolding protein binding site. L245 also binds R101 in the coat protein P-domain as well as E18 in the N-arm. These results demonstrate the presence of secondary coat binding sites that may function in conjunction with the HTH domain to promote the assembly of procapsids with the correct curvature.

**Importance:** Many dsDNA viruses, including tailed bacteriophages and Herpesviruses, assemble precursor capsids, or procapsids, using an essential catalytic scaffolding protein. How scaffolding proteins induce proper assembly of their major capsid proteins remains unclear. The scaffolding protein of bacteriophage P22 has a C-terminal helix-turn-helix domain that interacts with the N-arm of the coat protein to activate it for assembly. Here, a second potential coat protein interaction site is identified in scaffolding protein at residue L245. Residue L245 may be involved in stabilizing a small helical domain based the effect of the substitution scaffolding protein stability and a predicted model of the scaffolding protein fold, thereby indicating additional complexity in the interaction between coat and scaffolding proteins.

## Introduction

Across many double-stranded (ds) DNA bacteriophages as well as herpesviruses and adenoviruses, scaffolding proteins act as catalysts to drive procapsid assembly (1-3). Bacteriophage P22, which infects *Salmonella enterica* serovar Typhimurium, has been studied extensively since the 1950s and the functional role of its scaffolding protein (gp8) in assembly has been particularly well characterized (1, 3). In P22, other tailed bacteriophages and some dsDNA viruses, scaffolding protein interacts with structural proteins which can include coat, portal, and ejection proteins, to ensure procapsids are the correct size and contain the required components to mature into virions (2, 4-8). During P22 morphogenesis, scaffolding protein initiates procapsid assembly by driving the oligomerization of the dodecameric portal ring, which ensures portal incorporation at a single vertex (9-11). Scaffolding protein co-assembles with coat protein, nucleating at the portal complex, to form a shell of coat protein with internally located scaffolding protein (4, 12, 13). DNA is packaged through the portal vertex and scaffolding protein is released through holes in the procapsid to be recycled for further rounds of assembly (2). Once DNA is packaged, plug proteins and the tail machinery are added to complete mature virions (11). Assembly in the absence of scaffolding protein results in the formation of aberrant particles as well as empty T=7 procapsid-like particles and T=4 petite particles that cannot package DNA (14-16). Thus, scaffolding protein is required for the efficient nucleation and assembly of procapsids that are maturation competent.

Scaffolding proteins are generally shown to have intrinsically disordered regions (IDRs). The 303 amino acid P22 scaffolding protein is also disordered, due to its charge distribution, flexibility, and the apparent absence of a well-formed hydrophobic core (17-20). These characteristics enable scaffolding protein to interact with its numerous protein binding partners but make structural characterization challenging. In solution, scaffolding protein is an elongated α-helical protein that exists primarily as monomers and dimers, as well as tetramers at high concentrations (18, 21, 22). CryoEM reconstructions of P22 procapsids show only small regions of density attributed to scaffolding protein bound to coat protein hexamers around the three-fold axes of symmetry, indicating most of the scaffolding protein is disordered even when bound to procapsids (13, 23-25). The density likely corresponds to the C-terminal coat binding domain, which forms a helix-turn-helix (HTH) structure (26).

As the only known structured region of scaffolding protein, the HTH has been subjected to rigorous biochemical analysis. The minimal coat binding domain is comprised of residues 280-294. Deletion of the C-terminal 11 residues and other disruptions of the of the HTH structure result in a complete loss of function for procapsid assembly (24, 27, 28). Coat:scaffolding protein bindings occur via electrostatic interactions (29, 30). The HTH domain has a strong positive charge and residue R293 is essential for interaction with D14 on the coat protein N-arm (31-34). The coat protein N-arm and P-domain undergo significant structural rearrangements during maturation, which likely facilitates scaffolding protein release (23, 33, 34). The coat protein R101C substitution located at the amino-terminus of the spine helix in the P-domain also impedes scaffolding protein interactions (33, 34). Many other mutants support the hypothesis that coat:scaffolding protein interactions must be tightly regulated in order to support procapsid assembly and maturation (35).

While the C-terminal HTH of scaffolding protein is involved in coat binding, other regions have additional functional roles. The N-terminus is implicated in translational autoregulation and exit from the procapsid, as well as incorporation of the ejection proteins (28, 36-38). Truncation of the N-terminal 57 scaffolding protein residues causes incomplete DNA packaging due to retention of ∼140 molecules of scaffolding protein within the procapsid (28). This mutant and the temperature-(*ts*) and cold-sensitive (*cs*) *ts/cs*Q149W and *ts*L177I mutants fail to package the ejection protein, gp16, and are unable to undergo normal DNA packaging (5, 28). A region involving residues 229-238 is implicated in incorporation of portal protein (28). However, mutants S242F and Y214W, which lie outside the putative portal binding region, also fail to incorporate portal (5, 28). In total, these data indicate that scaffolding protein may have a more complex tertiary structure than previously suggested.

## Methods

### Phage and procapsid production

The host for bacteriophage P22 replication was *Salmonella enterica* serovar Typhimurium DB7136 (*leuA414*^*-*^*am hisC525*^*-*^*am sup*^*0*^). A su^+^ derivative strain of DB7136, *Salmonella enterica* serovar Typhimurium DB7155 (*supE20 leuA414*^*-*^*am hisC525*^*-*^*am*), was used for propagating virions with amber mutations (39). For complementation experiments, two P22 strains that are unable to express scaffolding protein (gene 8) due to an amber mutation, were used. To inhibit lysogeny, both strains contained the c1-7allele. P22 8^-^*amH202* c1-7 was used for efficiency of plating (EOP) experiments. P22 8^-^*amH202* 13^-^amH101 c1-7, which contains an amber mutation in gene 13 to prevent host cell lysis, was used experiments to produce procapsids and virions *in vivo* (4).

### Untagged scaffolding protein vectors and strains

Gene 8 was cloned into pASG-IBA105 (IBA Lifesciences, Götingen, Germany) using the direct cloning method described in the StarGate® direct transfer cloning protocol (IBA Lifesciences) with restriction enzyme Esp3I, to generate the plasmid pCLD01. This plasmid is inducible with 30 μg/L AHT anhydrotetracycline (AHT) and encodes gene 8. The N-terminal Twin Strep-tag was deleted via inverse polymerase chain reaction (PCR) using non-overlapping primers flanking the tag, resulting in plasmid pCLD02 (40). Site-directed mutagenesis using non-overlapping primers and inverse PCR was also used to introduce the L245A mutation in the scaffolding protein gene. PCR reactions, prepared in Phusion high-fidelity master mix with high-fidelity buffer (NEB), were run and then phosphorylated and re-ligated using T4 polynucleotide kinase (NEB) and T4 ligase (NEB, Ipswich, MA) respectively. The constructs were verified by DNA sequencing prior to transformation by electroporation into DB7136 to be utilized in complementation experiments. The plasmid confers ampicillin resistance.

### His_6_-tagged scaffolding protein vectors and strains

Gene 8 was synthesized and cloned into pet45b using the KpnI and BamHI sites by GenScript (Piscataway, NJ). The plasmids containing the L245A and L245C substitutions in gene 8 were also made by GenScript. The constructs, which confer ampicillin resistance, were chemically transformed into *Escherichia coli* BL21 (DE3) cells to express his_6_-tagged scaffolding protein under the control of a T7 promotor with 0.1 mM isopropyl β-D-1-thiogalactopyranoside (IPTG).

#### Efficiency of Plating (EOP) assay

*Salmonella* DB7136 cells containing the plasmid pCLD02 encoding WT or mutated gene 8 were grown to mid-log phase (∼2 × 10^8^ cell/mL) at 30 °C prior to being harvested by centrifugation at 4000 x g for 10 mins at 4°C. The cells were resuspended in 1/10^th^ of the original culture volume with ice-cold Luria-Bertani (LB) broth. Phages containing the 8^-^*am* allele were serially diluted in dilution fluid (10 mM Tris, pH7.6, 100 mM MgCl_2_) and added to soft agar, followed by cells and inducer (30 μg/L AHT). The mixture was plated on LB agar with 100 μg/mL ampicillin and incubated overnight at temperatures from 22 - 41 °C or for two nights at 16 °C before counting the plaques. Relative titer was calculated by dividing the titer of the 8^-^ *am* infections at the varied temperatures complemented by expression of WT or mutated gene 8 in pCLD02 by that of the expression of WT scaffolding gene at 30°C.

#### *In vivo* production of procapsids and virions

*Salmonella* DB7136 cells containing the plasmid pCLD02 encoding WT or mutated gene 8 were grown at 22 and 39°C to mid-log phase before being induced with 30 μg/L AHT and infected with 8^-^*am* 13^-^*am* P22 at a multiplicity of infection (MOI) of 5. Growth continued at their respective temperatures for 4 hours. Cells were harvested by centrifugation at 14,000 xg (Thermo, Fiberlite F12-6×500 LEX rotor) for 10 mins at 4°C and resuspended in ice-cold buffer (20 mM Tris pH 7.6, 100 mM MgCl_2_) prior to being stored at -20°C. Cells were lysed by freeze-thawing twice. Protease inhibitor cocktail (Roche), 1 mM phenylmethyl sulfonyl fluoride (PMSF), and 100 μg/mL DNase and RNase were added to the cells prior to removal of the cell debris by centrifugation at 12880 x g (Thermo, Fiberlite F20-12×50 LEX rotor) for 10 min at 4°C. The supernatant was then ultracentrifuged at 163,072 x g (Thermo, T-865 rotor) for 45 mins at 4°C to pellet the procapsids and phages. The pellet was gently resuspended in 2 mL ice-cold buffer (20 mM Tris pH 7.6, 100 mM MgCl_2_) overnight on a shaker table at 4 °C.

#### Cesium chloride step gradients

Cesium chloride density step gradients were used to separate procapids from virions. Solutions of 1.6 and 1.4 gm/cc CsCl and 25% sucrose were prepared in 20 mM HEPES pH 7.6, 100 mM MgCl_2_. Gradients were prepared by layering 1.6 gm/cc CsCl on the bottom with 1.4 gm/cc CsCl, followed by the 25% sucrose and finally the sample of resuspended procapsids and phages on top. Samples were then centrifuged at 70000 x g for 1hr at 18 °C. Procapsids and phages, which sediment at the boundaries between the sucrose cushion and 1.4 gm/cc CsCl and between the 1.4 gm/cc and 1.6 gm/cc respectively, were extracted and dialyzed against 1 L 20 mM Tris pH 7.6, 100 mM MgCl_2_ overnight.

#### Negative-stain Transmission Electron Microscopy (TEM)

Grids were prepared to visualize procapsids and phages using transmission electron microscopy. Procapsid samples from *in vivo* experiments were diluted 1:1 with 20 mM Tris pH 7.6, 100 mM MgCl_2_, those from *in vitro* assembly were not diluted. Samples were loaded onto 300 mesh carbon coated copper grids (Electron Microscopy Sciences, Hatfield, PA) and stained with 1% uranyl acetate (UA). For phage samples, grids were glow discharged in a plasma cleaner prior to staining with 1% UA. Images were taken at 68,000x magnification on an FEI Tecnai TEM.

### SDS-PAGE of procapsids and phages

Concentrations of procapsids (ε=0.772 (mg/mL)^-1^ cm^-1^) and phages (ε=0.957 (mg/mL)^-1^ cm^-1^) were quantified by diluting samples 1:40 in 6 M guanidine hydrochloride (GuHCl) and measuring the A_280_. Equal amounts of protein were loaded onto 12.5% acrylamide gels and run at 35 mA/gel. Gels were silver stained prior to being imaged on a ChemiDoc MP imaging system (BioRad).

### Purification of his_6_-tagged scaffolding proteins

BL21 cells expressing WT, L245A, and L245C scaffolding protein from pET45b were grown at 37°C to mid-log phase (OD_600_=0.5). After inducing with 0.1mM IPTG, cells continued to grow for 4 hours. Cells were pelleted at 8000 x g (Thermo, Fiberlite F9-6×1000 LEX) for 10 mins at 4°C prior to resuspension in binding buffer (20 mM sodium phosphate pH 7.6, 10 mM imidazole, 300 mM sodium chloride) and storage at -20°C. After thawing cells on ice, 1 mM PMSF, and 100 μg/mL DNase and RNase were added. Sonication was used to lyse the cells and debris was pelleted by centrifugation at 12880 x g (Thermo, Fiberlite F20-12×50 LEX rotor) for 10 mins at 4°C. Scaffolding protein was purified on a TALON Superflow metal affinity column (Takara Bio, San Jose, CA). Fractions containing scaffolding protein were pooled and concentrated in 12-14kDa MWCO dialysis tubing (Spectrum Chemical, New Brunswick, NJ) by dehydration with PEG 20,000K to a concentration >5mg/mL. The concentration of the purified scaffolding protein was monitored by Pierce 660 Assay (ThermoFisher Scientific, Waltham, MA). The protein was dialyzed against 1 L of 25 mM Tris, 50 mM NaCl, pH 7.6 at overnight at 4 °C, the concentration was measured again, and aliquots were stored at -80 °C.

### Circular Dichroism

Prior to performing circular dichroism (CD), scaffolding proteins (his_6_-WT, his_6_-L245A, and his_6_-L245C) were dialyzed against 10 mM phosphate (pH 7.6) buffer at 4 °C overnight. Scaffolding protein concentrations were checked by Pierce 660 assay and samples were diluted using the dialysis buffer to 0.2 mg/mL. Samples were loaded into a 0.1 cm pathlength quartz cuvette with (Starna Cells Inc., Atascadero, CA) and CD spectra were obtained using a Chirascan V100 spectrometer (Applied Photophysics, Skipton, UK). Spectra were collected between 195-260 nm with 1 nm intervals, 1 nm bandwidth, and a time-per-point averaging of 5 sec at 20 °C. Thermal denaturation was done by increasing the temperature from 10 - 80°C in 2 °C steps at a ramp rate of 0.5 °C per minute with time-per-point averaging of 0.75 sec, bandwidth of 1 nm, and 1 nm intervals.

### *In vitro* procapsid-like particle (PLP) assembly

Coat monomers were produced from shells by denaturation with 9 M urea followed by dialysis with 3 × 1 L of 20 mM HEPES pH 7.6 buffer, repeated twice for 2-3 hours and once overnight. Assembly reactions were initiated by adding 0.5 mg/mL coat monomers to 0.5 mg/mL scaffolding protein in 20 mM HEPES, 70 mM potassium acetate (KAc) buffer, pH 7.6. Tris(2-carboxyethyl)phosphine hydrochloride (TCEP, Sigma-Aldrich), 5 mM, was added for the assembly reaction with his_6_-L245C scaffolding protein. Assembly was monitored in a Horiba Fluoromax-4 Fluorometer (Irvine, CA) by light scattering at 500 nm for 3000 sec at 20 °C. The assembly products were run on a 1% SeaKem LE agarose gel at 100 V for 1 hr and Coomassie stained. The gel was imaged on a ChemiDoc MP imaging system (BioRad). Reaction products were also visualized by TEM.

### Scaffolding protein reincorporation into empty procapsid shells

Empty procapsid shells were made using the method described in (41). Briefly, the procapsids were repeatedly incubated with 0.5 M guanidine hydrochloride, each time followed by centrifugation to pellet the shells and resuspension by gentle shaking with buffer, until there is little scaffolding protein remaining. Reincorporation, or “restuffing”, was done by incubating 0.2 mg/mL scaffolding protein (his_6_-WT, his_6_-L245A, and his_6_-L245C) with 0.33 mg/mL shells in 20 mM HEPES, 70 mM potassium acetate pH 7.6 buffer at 22 °C for 1 hr. Each reaction was applied to a linear 2 mL 5-20% sucrose gradient made with the same buffer. Gradients were centrifuged at 45000 x g (Thermo S120AT2 rotor) for 35 min at 20 °C before being fractionated into 100 μL aliquots. Gel samples were prepared with 10 μL of 5X sample buffer with β-mercaptoethanol (BME) and 40 μL of each fraction and used for SDS-PAGE on a 10% acrylamide gel. Gels were Coomassie stained and destained with 10% acetic acid prior to imaging on a ChemiDoc MP imaging system (BioRad).

#### Crosslinking L245C scaffolding protein with cysteine mutant shells

Restuffing reactions with 0.2 mg/mL his_6_-L245C scaffolding protein and 0.33 mg/mL shells with D163C or D198C coat protein were performed in 20 mM sodium phosphate buffer pH 7.6 with 5 mM TCEP for 1 hr at 22 °C. Control reactions with L245C scaffolding protein alone, and D163C or D198C shells alone were produced with the same conditions. The restuffing reactions were then incubated with 100 μL of 50% TALON slurry (TakaraBio, Kusatsu, Japan), previously washed with a ten-fold excess of 20 mM phosphate buffer three times, to remove unbound scaffolding protein. The TALON resin was pelleted by centrifugation at 14,000 x g, for 5 minutes at 4 °C and the supernatant was removed. The restuffing supernatant and control reactions were dialyzed in Pur-A-lyzer Midi 6000 dialysis cups (MilliporeSigma, Burlington MA) overnight against 500 mL of phosphate buffer at 4 °C to remove the TCEP. The sample volume was divided into thirds, one for a pre-crosslinking sample and two other samples for crosslinking reactions. Reactions were crosslinked with the non-cleavable crosslinkers 200 μM bismaleimidoethane (BMOE; 8 Å crosslinking arm) or 80 μM 1,11-bismaleimido-triethyleneglycol (BM(PEG)_3_; 14.7 Å crosslinking arm) for 1 hour at room temperature and quenched with 20 mM L-cysteine. SDS-PAGE was performed for all samples using 7.5% acrylamide gels. To assess the formation of crosslinks, gels were silver stained and imaged on a ChemiDoc MP imaging system (BioRad).

## Results

Due to its intrinsic disorder, a complete structure of P22 scaffolding protein has not been possible, making mapping the protein-protein interaction sites challenging. To identify scaffolding sites required for interacting with binding partners, the effects of alanine mutants generated at various positions on scaffolding protein were characterized. Scaffolding protein residue L245 was found to be essential for interaction with coat protein.

### Residue L245 is critical for interaction with coat monomers *in vivo*

To initially assess the effect of substitutions, efficiency of plating (EOP) experiments were performed by infecting host *Salmonella*, transformed with a plasmid carrying WT or mutant scaffolding protein, with *8*^*-*^*am* P22 phage. The infections were plated and grown at temperatures ranging from 16 - 41 °C to assess virion growth. Calculating the relative titer indicates how well the plasmid-encoded mutated gene 8 complements the 8^-^*am* phage, which lacks a functional scaffolding protein gene.

L245A scaffolding protein caused a severe cold sensitive (*cs*) phenotype in EOP experiments. The relative titer was two orders of magnitude lower than WT even at the most permissive temperatures (37 - 41 °C), and at 16 °C, it was reduced to the reversion frequency of the 8^-^*am* phage (Fig. 1). This phenotype indicates that mutating scaffolding protein residue L245 affects virus morphogenesis in some way. To determine if the L245A substitution affects procapsid assembly or mature virion formation *in vivo*, particles from complementation experiments performed at both permissive and non-permissive temperatures (22 and 39 °C, respectively) were analyzed by negative stain transmission electron microscope (TEM) after separation by CsCl density gradient. At both temperatures, complementation of the 8^-^*am* phage with WT gene 8 resulted in the formation of T=7 procapsids and viral particles (Fig. 2A). Electron micrographs of particles formed at both temperatures with L245A scaffolding protein show normal sized (T=7) procapsids, as well as an abundance of aberrant spirals and petite (T=4) capsids, similar to that seen in an 8^-^*am* infection, indicating a defect in coat polymerization (14). No phages were detected on the TEM grids from particles isolated from the complementation experiment where gene 8 carrying the L245A mutation was expressed at 22 °C, described above. However, in the mature virion samples from complementation at 39 °C, some mature, DNA-filled, tailed phages were detected, consistent with the EOP experiments (Fig. 2A).

**FIG 1:**
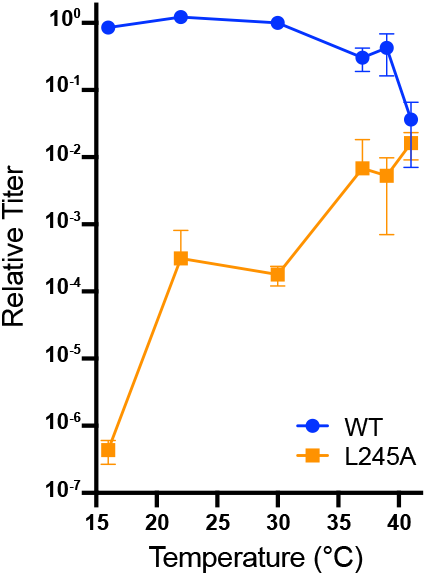
The L245A substitution in scaffolding protein confers a cold-sensitive (*cs*) phenotype in complementation experiments. Efficiency of plating (EOP) shows the average titer of phages assembled using WT (blue) or L245A (orange) scaffolding protein at 16, 22, 30, 37, 39, and 41 °C, relative to those produced with WT scaffolding protein at 30 °C.

**FIG 2:**
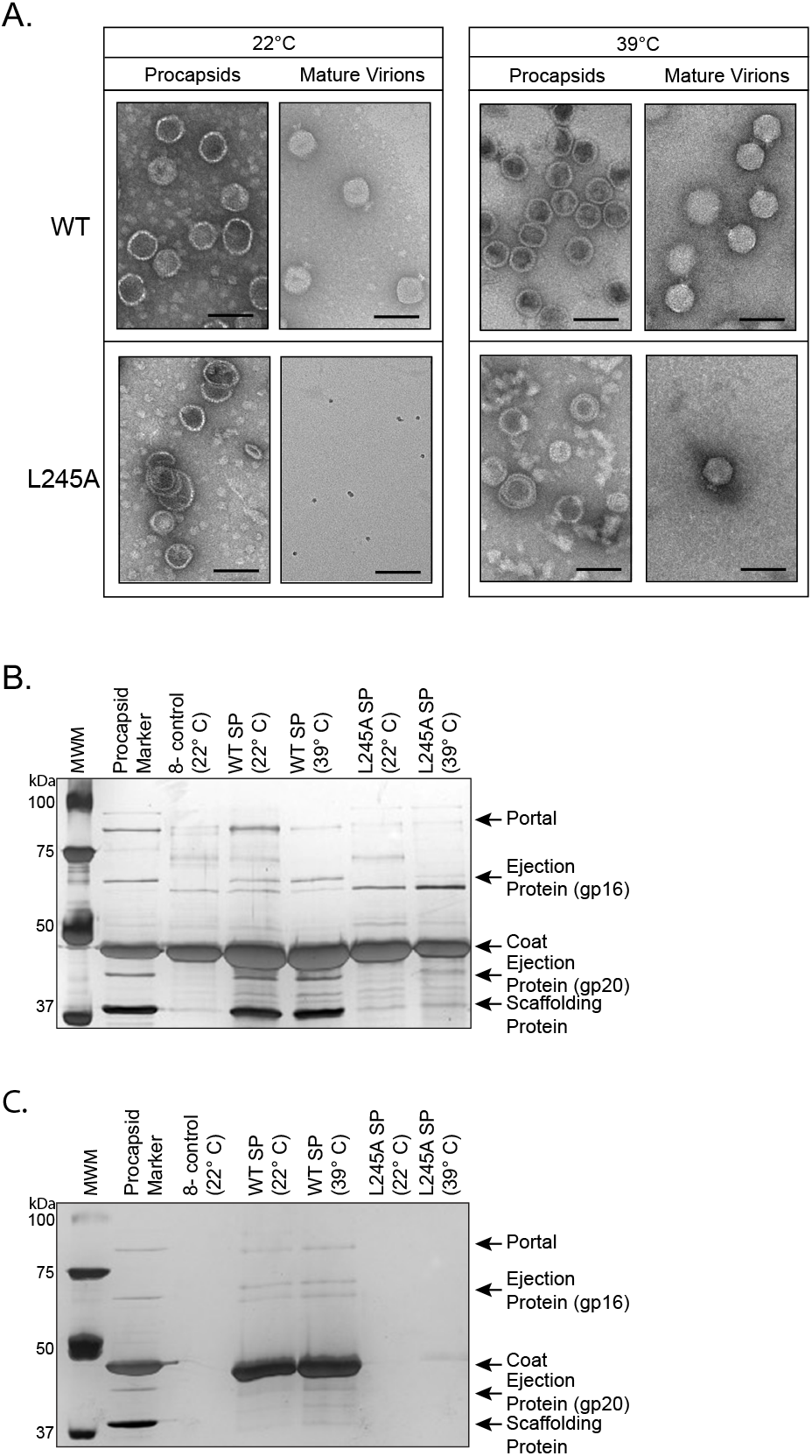
*In vivo* assembly with L245A scaffolding protein yields aberrant particles. Aberrant particles, procapsids and phages are produced from *in vivo* complementation experiments at 22 and 39 °C with WT and L245A scaffolding protein. Particles were separated using cesium chloride step gradients. (A) Negative stain transmission electron micrographs. Scale bars are 100 nm. Silver stained 12.5% SDS acrylamide gels of the (B) aberrant particles, procapsids and (C) phages shown in (A).

The effects of L245A scaffolding protein on the incorporation of the portal complex and ejection proteins into the procapsid were evaluated as scaffolding protein is required for on these functions, as well (Fig. 2B,C). Procapsid and mature virion samples were analyzed by SDS-PAGE. In procapsids and phages assembled with WT scaffolding protein at 22 or 39 °C, portal protein and the ejection proteins gp20 and gp16 were present (ejection protein gp7 is not retained on these gels), indicating that scaffolding protein was functioning properly. Conversely, when scaffolding protein L245A was used in the complementation experiments, the minor proteins were not present in particles assembled at 22 °C, due to the absence of scaffolding protein (Fig. 2B,C). (4, 14). However, both gp16 and gp20 are visible in particles produced with L245A scaffolding protein at 39 °C, indicating that they are packaged into procapsids but with decreased efficiency. Portal and ejection proteins were not detected in the gels of the L245A phage samples, due to the low abundance of mature virions (Fig. 2C). Together, these data indicate that residue L245 is important for proper interaction with coat protein, so the lack of portal and ejection proteins was likely due to the low concentration of scaffolding protein in the particles.

### L245A scaffolding protein interacts weakly with coat monomers and procapsids *in vitro*

Based on our *in vivo* experiments, we hypothesized that L245 is a coat protein interaction site. To test this hypothesis, we performed *in vitro* assembly reactions and affinity chromatography with purified coat monomers and his_6_-tagged scaffolding proteins. The results of the *in vitro* assembly reactions, where purified coat protein monomers are mixed with purified his_6_-tagged scaffolding protein, showed that procapsid assembly proceeds slowly for L245A scaffolding protein compared to WT scaffolding protein, though more efficiently than coat protein assembled in the absence of scaffolding protein (Fig. 3A). His_6_-tagged L245C scaffolding protein, which was used for crosslinking experiments (below), was as efficient at catalyzing assembly as the his_6_-tagged WT scaffolding protein, indicating that the cys substitution does not affect interaction with coat protein.

**FIG 3:**
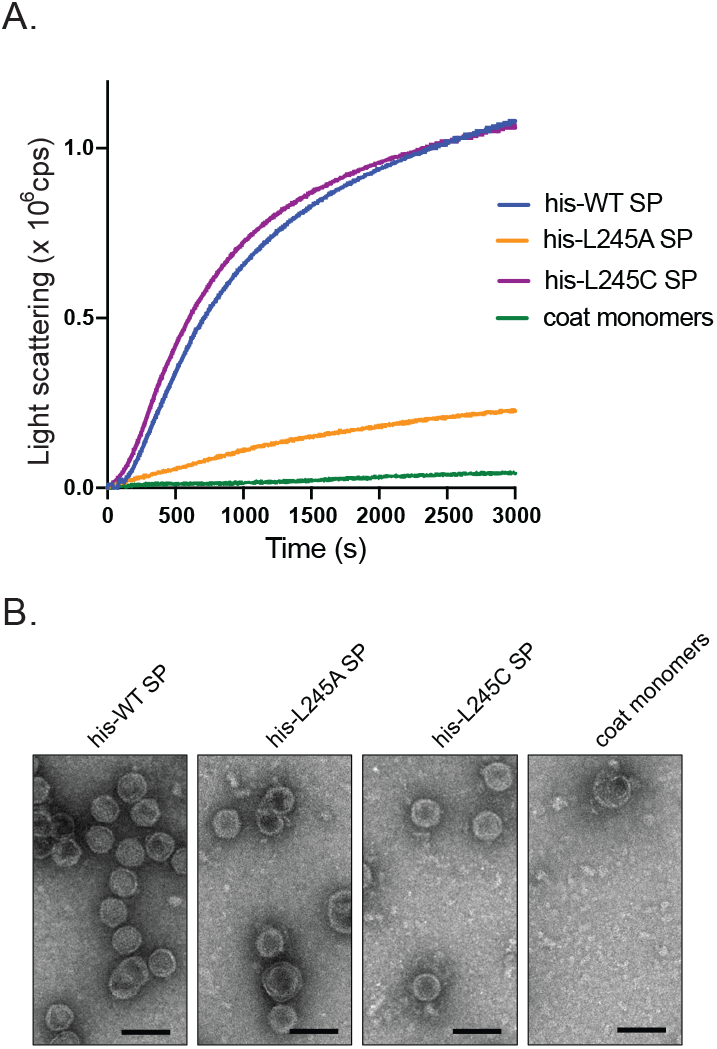
*In vitro* assembly of procapsids with mutant scaffolding proteins. (A) Kinetics of *in vitro* procapsid assembly of coat monomers with his_6_-WT scaffolding protein or scaffolding protein mutants his_6_-L245A and his_6_-L245C measured by light scattering at 500 nm. (B) Negative-stained TEM images of *in vitro* assembly reactions. Scale bars are 100 nm.

The products of the assembly reactions were imaged by negative-stain TEM to assess the fidelity of procapsid assembly with the different scaffolding protein mutants (Fig. 3B). While his_6_-tagged WT and L245C scaffolding proteins assembled normal procapsids, the majority of the particles produced by his_6_-tagged L245A scaffolding protein were aberrant or petites. Coat monomers alone acted as a negative control; the products of this reaction were a few aberrant spirals and petite particles. These results are consistent with the light scattering data, and support our hypothesis that L245A scaffolding protein does not interact properly with coat protein, as this is a result that is similar to assembly without scaffolding protein *in vivo* and *in vitro* (5, 7).

These results were further corroborated by weak affinity chromatography. Coat monomers and ovalbumin (a negative control) were applied to metal-affinity columns bound with his_6_-tagged WT or his_6_-tagged L245A scaffolding protein and elution was monitored by tryptophan fluorescence (Fig. 4). The retention of coat protein on the column is qualitatively related to the strength of its binding to WT or mutant scaffolding protein (33, 42, 43). The ovalbumin eluted with a peak centered at 2.1 mL from columns generated with either his_6_-tagged WT and L245A scaffolding proteins, indicating the volume at which a non-interacting protein elutes. The coat protein elution profiles for both the his_6_-tagged WT and his_6_-tagged L245A scaffolding proteins showed a doublet of peaks at ∼ 2.4 and 4.2 mL. This profile is consistent with previous studies showing that coat protein monomers sometimes have a fraction that may not interact well with scaffolding protein attached to a column (32). A greater fraction of coat protein eluted from the WT scaffolding protein column in the later peak (4.2 mL) rather than the early eluting peak (2.4 mL) (Fig. 4A), indicating that a significant portion of the coat protein was able to interact with WT scaffolding protein. In contrast, the coat protein elution profile from the L245A scaffolding protein column shows a higher initial peak at 2.4 mL followed by broader peak centered at 4.2 mL. The differences in the elution profiles indicate that the L245A scaffolding protein substitution weakens binding to coat protein. Together, these *in vitro* results suggest that disruption at site L245 likely affects a coat interaction site.

**FIG 4:**
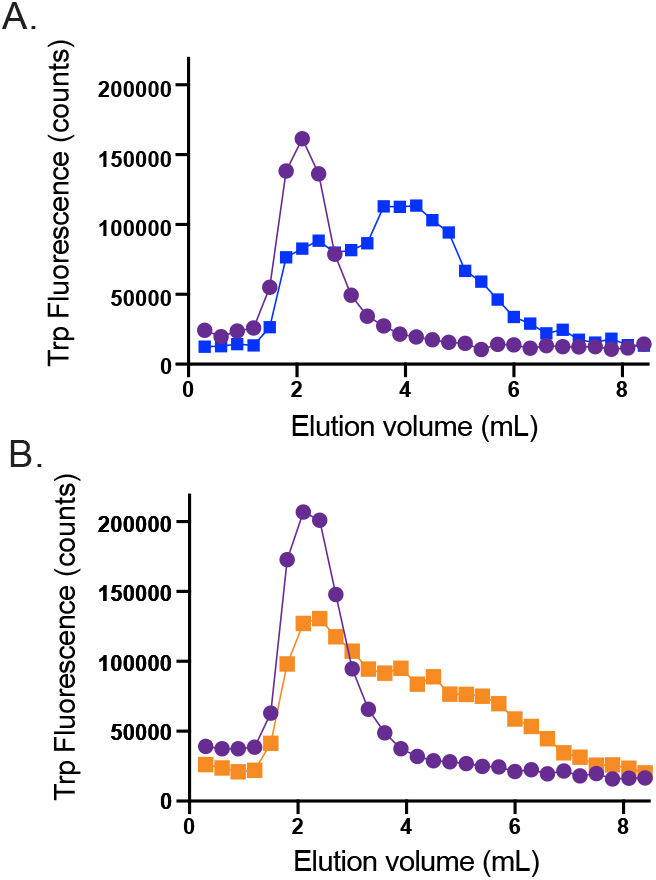
Coat protein binds more weakly to L245A scaffolding protein. Weak affinity chromatography with a (A) his_6_-tagged WT scaffolding protein column or (B) his_6_-tagged L245A scaffolding protein column. Elution was measured by tryptophan fluorescence (cps) and normalized to the number of tryptophan residues in each protein. The elution of ovalbumin is indicated with purple circles, and the elution of WT coat protein monomers by blue or orange squares.

In addition to interacting with coat monomers to assemble procapsids, scaffolding protein has also been shown to repackage, or ‘restuff’, into procapsids that have been stripped of their scaffolding protein, known as shells (12). To understand whether the L245A mutation affects only interactions with monomers or if it also altered interactions with the procapsid shell interior, restuffing experiments were done by incubating his_6_-tagged WT, L245A, and L245C scaffolding protein with shells and running the reactions on 5-20% sucrose gradients to separate the unbound scaffolding protein from the restuffed procapsids. The gradients were fractionated and analyzed by SDS-PAGE on 10% acrylamide gels (Fig. 5). The his_6_-tagged WT, and his_6_-tagged L245C scaffolding protein can repackage into shells while L245A scaffolding protein does not efficiently restuff, suggesting that L245A scaffolding protein also interacts more weakly with assembled coat protein than the WT scaffolding protein.

**FIG 5:**
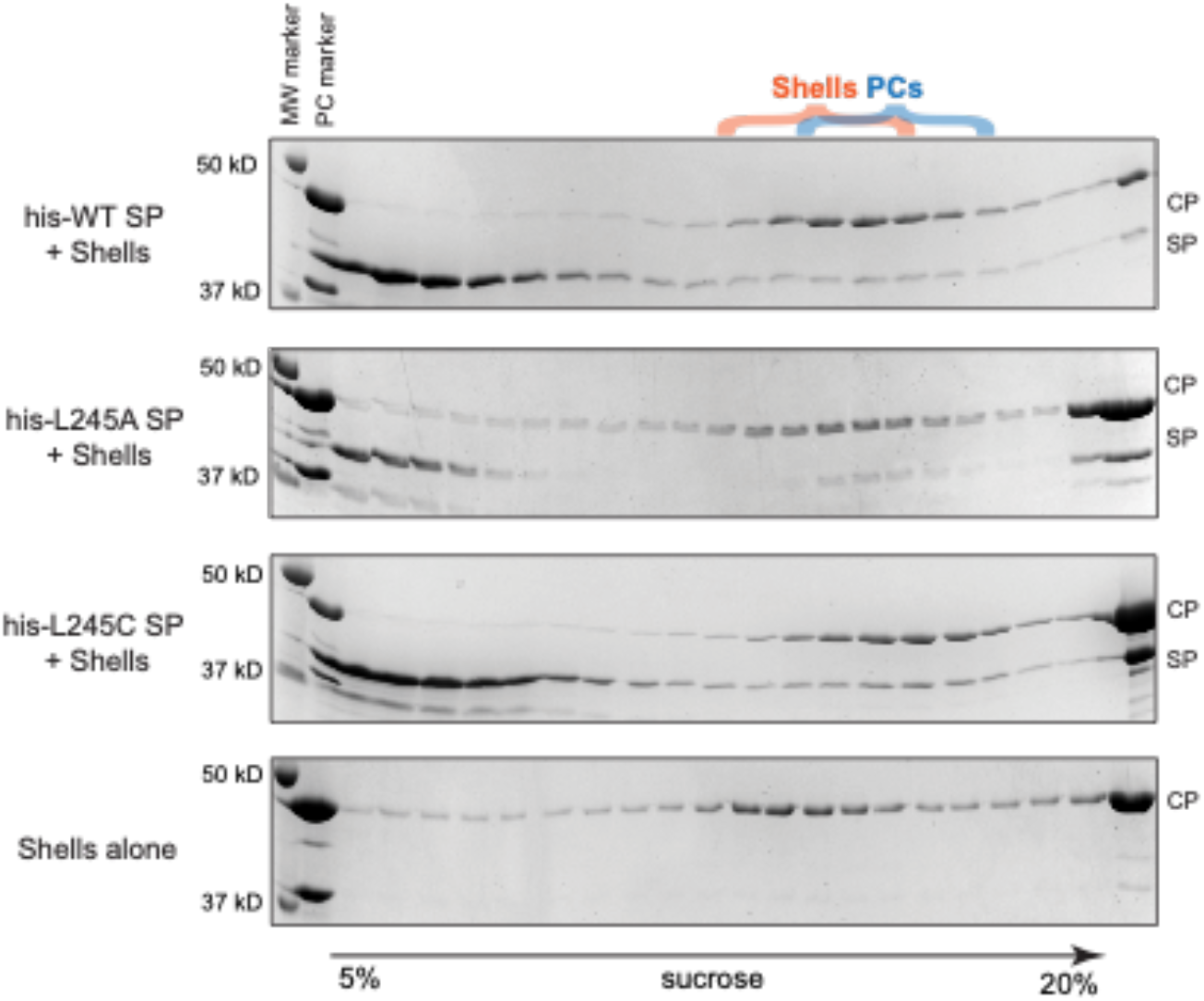
L245A scaffolding protein is less able to re-enter shells. SDS-PAGE of 5-20% sucrose gradient fractions from scaffolding protein restuffing reactions using P22 empty procapsids shells with his_6_-WT, his_6_-L245A, and his_6_-L245C scaffolding protein (SP). Coat protein (CP) and scaffolding Protein (SP) are denoted. The shell and procapsid peaks are indicated by orange and blue brackets respectively.

### L245A disrupts scaffolding protein secondary structure

Circular dichroism (CD) was performed to assess if secondary structural changes caused the coat binding defect of L245A scaffolding protein. CD spectra were taken for his_6_-tagged WT and L245 scaffolding proteins. At 10 °C, the proteins share essentially identical α-helicity that is characteristic of scaffolding protein, demonstrated by the strong negative peaks at 208 and 222 nm (Fig. 6A). As L245A causes a temperature-sensitive phenotype, thermal melts monitored with CD were performed to discern any differences in the unfolding and stability of L245A vs. WT scaffolding proteins. Scaffolding protein has a broad melting transition consistent with it having IDRs (44). Starting the melt at 10 °C (measured inside the cuvette) allowed observation of an early transition not previously appreciated in the WT scaffolding protein melt, seen as a plateau between ∼ 25 and 32 °C (Fig. 6B). This plateau was destabilized in the melt of L245A scaffolding protein, suggesting that the L245A substitution disrupted some rather unstable structure. The spectra of WT and L245A scaffolding proteins were taken at 30 °C, which showed that L245A scaffolding protein lost some negative ellipticity at 222 nm and at 192 nm (Fig. 6C). This result is consistent with diminished α-helical and increased irregular/disordered secondary structure as a result of the L245A substitution (45). Overall, these data suggest that the substitution L245A affects the secondary structure of scaffolding protein.

**FIG 6:**
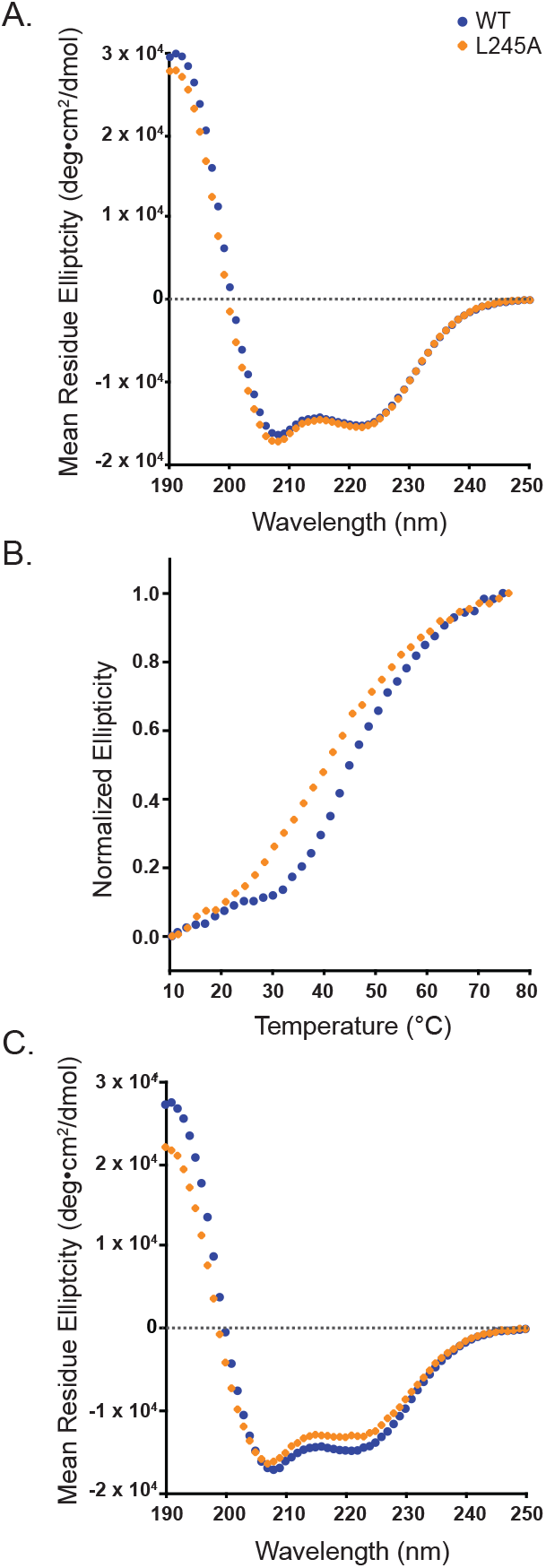
L245A scaffolding protein is less stable that WT scaffolding protein. (A) Circular dichroism spectra of WT and L244A scaffolding proteins taken at 10 °C. (B) Thermal melts showing that L245A has a thermal denaturation shifted to lower temperatures. (C) Spectra taken at 30 °C.

### Scaffolding protein residue L245 interacts with the N-arm and spine-helix of coat protein

To determine where the scaffolding protein residue L245 interacts with P22 coat protein, we performed crosslinking experiments with procapsid shells formed from coat protein cysteine mutants, restuffed with his_6_-tagged L245C scaffolding protein (Fig. 7). We chose coat protein mutants in the N-arm (E15C and E18C), the P-domain (R101C in the spine helix), as well as the A-domain (D163C and D198C), as these domains undergo significant rearrangement during maturation (23). N414C in the final anti-parallel β-strand of the P-domain is not predicted to move during maturation and serves as a negative control. Each of these mutants retains the native cysteine at position 405, which is not accessible to crosslinkers when assembled into shells (see WT shells, Fig. 7C) (6, 34). The N-arm residue D14 is the interaction site for residue R293 of the scaffolding protein HTH domain, while the A-domain has been linked to scaffolding protein release as it unfurls to close the exit holes in the hexons (23, 32, 46). The P-domain has also been shown to undergo significant conformational changes during maturation, especially in the P-loop (23, 46). R101C is a *ts* coat substitution that was previously shown to interact only weakly with WT scaffolding protein (33).

**FIG 7.**
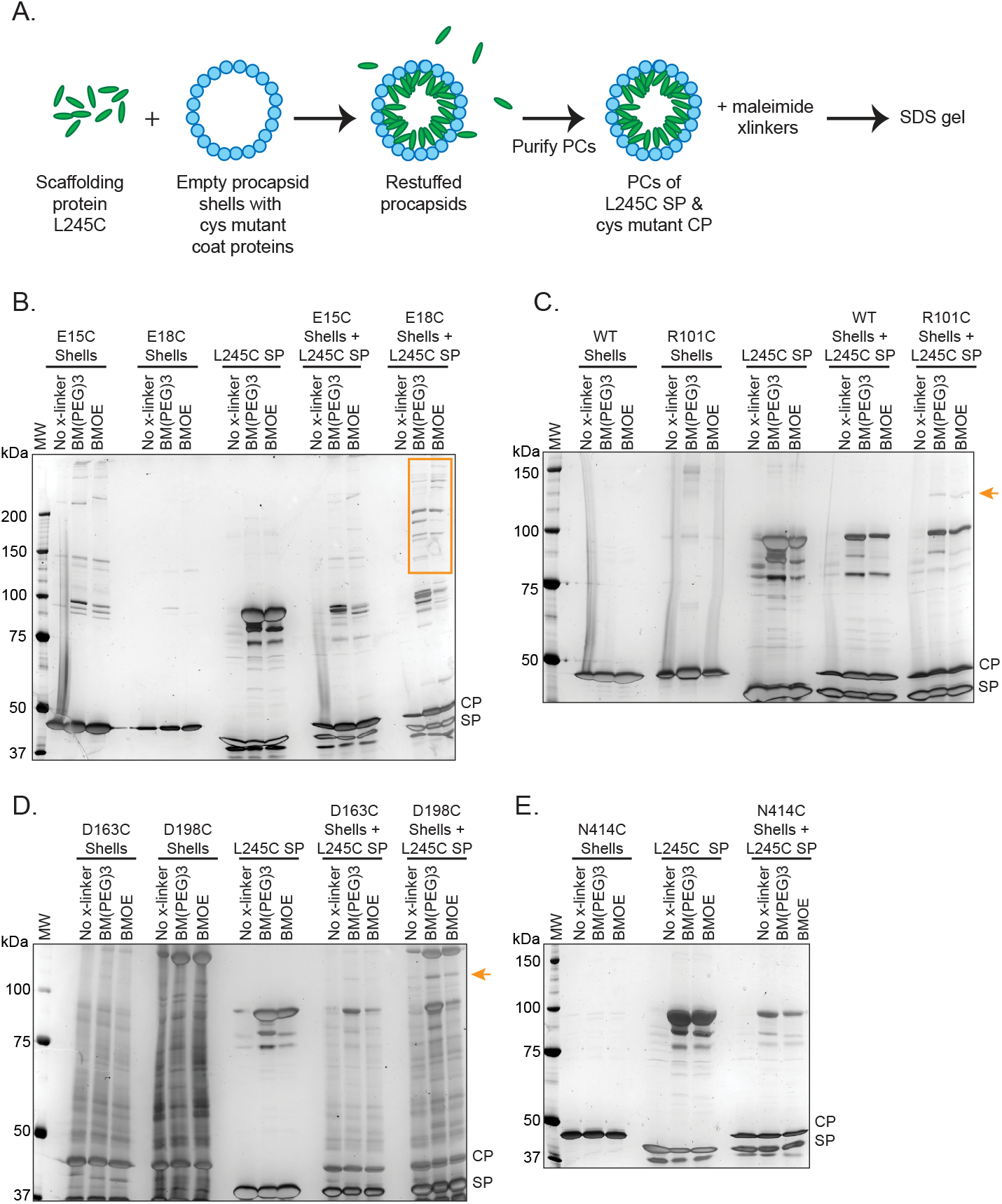
(A) Schematic showing the process of restuffing and crosslinking scaffolding protein (SP) his_6_-L245C with cysteine mutant empty coat protein shells. SDS-PAGE of the crosslinking reactions using BM(PEG)_3_ and BMOE with (B) E15C and E18C shells, (C) WT and R101C shells, (D) D163C and D198C shells, and (E) N414C shells. Crosslinked bands have been indicated with orange arrows and boxes.

Cysteine mutant shells, L245C scaffolding protein, or the L245C scaffolding protein restuffed into the mutant shells were treated with two maleimide crosslinking reagents with different crosslinking arm lengths, 1,11-bismaleimido-triethyleneglycol (BM(PEG)_3_, 14.7 Å), and bismaleimidoethane (BMOE, 8.0 Å). Each reaction was analyzed by 7.5% SDS-PAGE (Fig. 7B). Higher molecular weight bands were detected in the reactions with L245C scaffolding protein alone, indicating that this scaffolding protein mutant can form crosslinked dimers. Several crosslinks were detected between L245C scaffolding protein and the E18C coat protein (Fig. 7B), while a distinct band indicated L245C crosslinked to R101C (Fig. 7C) and D198C (Fig. 7D) coat protein shells using both using BM(PEG)_3_ and BMOE. No crosslinking was detected with N414C shells (Fig. 7E). These data indicate that scaffolding protein residue L245 is in contact with coat protein residues in the N-arm, P-domain and A-domain, which are sites consistent with data from P22 and other phages (see discussion).

## Discussion

Among the tailed, dsDNA bacteriophages, scaffolding proteins catalyze the assembly procapsids with the correct morphology and provide stability prior to maturation (1, 3, 6, 35, 47-51). The bacteriophage P22 scaffolding protein is among the best studied and its coat binding function has been extensively characterized. Coat-scaffolding interactions must be tightly controlled, as binding that is either too weak or too tight has detrimental effects on procapsid assembly (33, 35, 52, 53). In this study, we identify a novel site critical for coat-binding on the P22 scaffolding protein and determine interaction sites on the coat protein by crosslinking.

### The L245A substitution impairs the coat assembly function of the HTH

The scaffolding protein C-terminal HTH domain contains the minimal coat binding domain comprised of residues 280-294, which can polymerize coat protein into particles, albeit inefficiently (26-28, 31). Surprisingly the scaffolding protein mutant L245A, which causes a *cs* phenotype but lays outside the previously identified coat binding region, was shown to have a detrimental effect on coat protein binding and particle assembly *in vitro* and *in vivo*. These data suggest that L245A may either 1) be a secondary coat binding site in scaffolding protein, or 2) affect the ability of the C-terminal HTH domain to function. Although scaffolding protein lacks a single, unified hydrophobic core, a local hydrophobic zipper has been shown to stabilize the HTH domain, maintaining the correct orientation for coat-binding (24). Interestingly, the amino acid sequence surrounding L245 has an abundance of hydrophobic residues punctuated by charged arginine, lysine, and glutamic acid residues. L245 may stabilize some local hydrophobic core, similar to the hydrophobic zipper in the HTH domain (24), thereby allowing for proper presentation of charged residues to coat protein for assembly.

### A predicted model for scaffolding protein

To explore the above hypothesis, AlphaFold2 was used to generate a model of scaffolding protein (Fig. 8) (54, 55). The program predicts with good confidence an elongated, highly α-helical model, with a disordered N-terminus, an elongated helix (hatchet handle), and a C-terminal HTH-domain, which is consistent with the previous structural data (17, 18, 21, 26, 56). Also predicted is a 5-helix domain (denoted as the “hatchet head”) adjacent to the long helix of the “hatchet handle”, boxed in Fig. 8A. Previous crosslinking and FRET data indicate that scaffolding protein residue D18 is in close proximity to both the C-terminal residue R303 and residue D181 (located at the C-terminus of the helical hatchet handle in the model) (56). In contrast, D181C and R303C do not form an intramolecular disulfide bond. These results can be consistent with the AlphaFold2 model owing to the flexibility of the N-terminal unstructured region and the loop region connecting the HTH to the 5-helix domain (28, 56).

**FIG 8:**
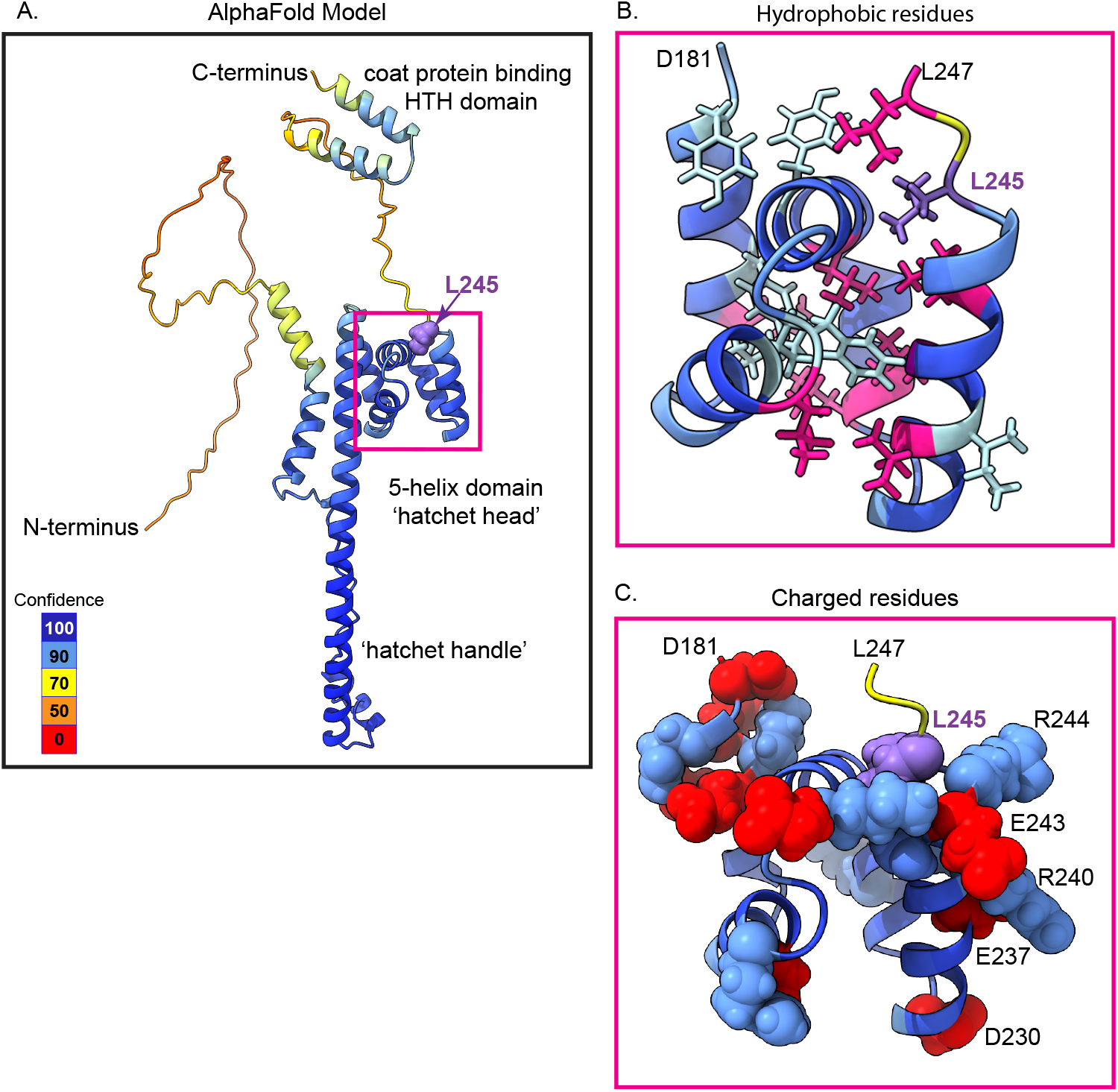
An AlphaFold2 prediction of the P22 scaffolding protein structure. (A) The entire scaffolding protein predicted structure, colored by confidence level. A magenta box shows the helical domain that contains residue L245 (purple spheres). (B) A zoom-in on the well-packed hydrophobic core of the 5-helix domain. L245 is indicated by purple sticks, all the other leucine residues are in magenta, and remaining hydrophobic residues are in light blue (isoleucines, valines, tyrosines and phenylalanines). (C) A zoom-in on the 5-helix domain to show the highly charged surface. Basic residues are indicated by blue spheres, acid residues in red spheres. The charged residues on the C-terminal helix are indicated by residue number. L245 is shown in purple spheres.

A scaffolding protein fragment comprised of residues 141-303 is fully capable of supporting assembly of procapsid-like particles *in vivo* and *in vitro* (21, 28). The 141-303 fragment has about half of the predicted hatchet handle helix plus the hatchet head and HTH domain. In contrast the scaffolding fragment containing residues 238-303, which crops the five helix hatchet head domain and the entire handle, can promote coat protein assembly but the particles produced are abnormal large spirals and aggregates (26). Thus, the hatchet head is likely important for regulating proper assembly of coat protein. L245 is modeled at the end of the C-terminal helix in the 5-helix bundle and proximal to residues L212 and L241, L247 (all the leucines are magenta in Fig. 8B), suggesting that it may be involved in stabilizing the hydrophobic core of the domain, consistent with our melt data showing decreased stability of the L245A mutant. While the alanine substitution may be too small to maintain these contacts, the L245C substitution does not impede coat protein interactions. The surface of the 5-helix hatchet head domain is very charged. Residues R244, E243, R240, E237 and D230 jut outwards (Fig. 8C), potentially providing a surface for electrostatic interactions with the coat protein or the scaffolding protein C-terminal HTH domain.

Due to to scaffolding protein’s flexibility and high number of binding partners, its tertiary structure and long-distance contacts have been proposed to play a significant role in regulating interaction sites (18, 24, 28, 35, 56). While this structure is only a prediction, it aligns well with prior structural evidence and provides new opportunities to propose and test hypotheses related to scaffolding protein structure and its binding domains. Based on this model, previous data and the data presented here, there are two possible hypotheses for the effects of the scaffolding protein L245A mutant. First, the 5-helix domain may stabilize or orient the HTH domain through direct interaction of the domains, resulting in a large, charged surface to facilitate coat-binding, and the L245A substitution negatively affects this interaction. Or second, the L245A substitution could destabilize the hydrophobic core within the hatchet head thereby causing tertiary structural changes that somehow alter the functionality of the HTH domain. These hypotheses will be tested by mutational analysis of the hydrophobic core.

### Scaffolding protein interactions with procapsids

Interactions between encapsulated scaffolding protein and procapsids have been probed through both biochemical and structural means. CryoEM reconstructions of scaffolding-filled and empty procapsids, as well as mature virions indicate that protein may bind inside the capsid around the 3-fold axis of symmetry, known as the “trimer tip” region, which undergoes significant conformational change during the maturation process (13, 23, 25, 46). The P-domain, N-arm, and A-domain undergo structural rearrangements upon DNA packaging that have been proposed to be important for scaffolding protein release (23, 34, 46). Additionally, several scaffolding-binding sites on coat protein have been identified. Cortines et al. identified the coat binding residues, R293 and K296, in the scaffolding protein HTH that are in close contact with negatively charged coat protein residues D14 and to a lesser extent E15 and E18 in the N-arm (31, 32). Teschke and Fong characterized the *cs* coat mutants T10I, which is also on the N-arm, and R101C in the coat protein P-domain. These both cause coat assembly defects and do not retain scaffolding protein within particles. T10, D14, and R101 are all in the vicinity of the “trimer tip” region, further supporting the structural data described above (6, 12, 13, 23). Multiple kinetic studies of coat-scaffolding protein and scaffolding protein-procapsid interactions have indicated at least two distinct binding classes exist for the ∼300 scaffolding binding sites in the capsid, 60 of which are high affinity sites with a *K*_*d*_ of 0.3 μM (12, 35, 57). Finally, the low resolution of scaffolding protein density in cryoEM reconstructions of the P22 procapsid (23) makes it unclear which scaffolding protein region(s) is bound to the interior of procapsids, but additional scaffolding protein domains are possible (56). Taken together, these data suggest that additional scaffolding-interaction sites are yet to be identified in the procapsid.

The crosslinking data presented here show that scaffolding protein residue L245 is near E18 on the coat protein N-arm and R101 in the spine helix of the coat P-domain. These data may indicate that residue L245 is adjacent and perhaps stabilizing the HTH at the 3-fold axis of symmetry. Alternatively, nearby charged residues on 5-helix domain could interact directly with the coat protein. In addition to interacting near the “trimer tip” regions in the procapsid, L245C crosslinks with D198C, positioned at the tip of the coat-protein A-domain, which flips outward during maturation and has been suggested to facilitate scaffolding protein exit (23, 34).

### Comparisons with other scaffolding proteins

Commonalities in coat-scaffolding interactions have been noted across the double-stranded DNA bacteriophages, as well as the *Microviridae* and *Herpesviridae*, showing that scaffolding proteins drive assembly and stabilize the quasi-stable procapsids through similar mechanisms (1, 3, 58, 59). In fact, owing to their shared coat protein HK97-fold, the general structural similarity between scaffolding proteins (e.g. having IDRs) and electrostatic interactions with coat protein at the N-arm (or N-helix in T7), is a repeated theme in bacteriophages P22, β29, SPP1, and T7 (32, 60-63). In SPP1, scaffolding protein gp11 binds both hexons and pentons through the spine helix, creating a network of molecular non-covalent staples connecting capsomers (60). Similar to P22 scaffolding protein, the phage T7 scaffold, gp9, has been shown to bind the A-domain in addition to the N-helix (62, 63). The data presented here are consistent with these results and suggest additional structural complexity of scaffolding protein that needs exploration.

## Acknowlegements

We thank Dr.Juliana Cortines for critical reading of this manuscript. We also thank Drs. Juliana Cortines and Charles Bridges for technical assistance.

